# Seeing our hand or a tool during visually-guided actions: different effects on the somatosensory and visual cortices

**DOI:** 10.1101/2022.11.04.515184

**Authors:** Benjamin Mathieu, Antonin Abillama, Simon Moré, Catherine Mercier, Martin Simoneau, Jérémy Danna, Laurence Mouchnino, Jean Blouin

**Affiliations:** Laboratoire de Neurosciences Cognitives (LNC), Aix-Marseille Université/ CNRS. Marseille, France; Centre interdisciplinaire de recherche en réadaptation et intégration sociale (CIRRIS) du CIUSSS de la Capitale-Nationale, Québec, Québec, Canada; Faculté de médecine, Université Laval, Québec, Canada; Institut Universitaire de France (IUF), Paris, France

**Keywords:** Electroencephalography, Proprioception, Sensory conflict, Vision, Sensory gating, Body representation

## Abstract

The processing of proprioceptive information in the context of a conflict between visual and somatosensory feedbacks deteriorates motor performance. Previous studies have shown that seeing one’s hand increases the weighting assigned to arm somatosensory inputs. In this light, we hypothesized that the sensory conflict, when tracing the contour of a shape with mirror-reversed vision, will be greater for participants who trace with a stylus seen in their hand (Hand group, n=17) than for participants who trace with the tip of rod without seen their hand (Tool group, n=15). Based on this hypothesis, we predicted that the tracing performance with mirror vision will be more deteriorated for the Hand group than for the Tool group, and we predicted a greater gating of somatosensory information for the Hand group to reduce the sensory conflict. The participants of both groups followed the outline of a shape in two visual conditions. Direct vision: the participants saw the hand or portion of a light 40 cm rod directly. Mirror Vision: the hand or the rod was seen through a mirror. We measured tracing performance using a digitizing tablet and the cortical activity with electroencephalography. Behavioral analyses revealed that the tracing performance of both groups was similarly impaired by mirror vision. However, contrasting the spectral content of the cortical oscillatory activity between the Mirror and Direct conditions, we observed that tracing with mirror vision resulted in significantly larger alpha (8-12 Hz) and beta (15-25 Hz) powers in the somatosensory cortex for participants of the Hand group. The somatosensory alpha and beta powers did not significantly differ between Mirror and Direct vision conditions for the Tool group. For both groups, tracing with mirror vision altered the activity of the visual cortex: decreased alpha power for the Hand group, decreased alpha and beta power for the Tool group. Overall, these results suggest that seeing the hand enhanced the sensory conflict when tracing with mirror vision and that the increase of alpha and beta powers in the somatosensory cortex served to reduce the weight assigned to somatosensory information. The increased activity of the visual cortex observed for both groups in the mirror vision condition suggests greater visual processing with increased task difficulty. Finally, the fact that the participants of the Tool group did not show better tracing performance than those of the Hand group suggests that tracing deterioration resulted from a sensorimotor conflict (as opposed to a visuo-proprioceptive conflict).

## 1. Introduction

Hands and fingers can be moved with extraordinary precision, notably when interacting with the external world. To successfully control movements with high spatial constraints, the brain uses two main sources of feedback: visual and somatosensory. Although these feedbacks first reach highly sensory-specific areas of the brain (e.g., the primary visual and somatosensory areas), they rapidly converge at common integrative areas (e.g., posterior parietal cortex; see Murray & Wallace, 2012 for a review). Importantly, the great adaptability of the sensorimotor system enables visual and somatosensory information to be spatially (and temporally) congruent. In order words, we see our hand where we feel it, and we feel our hand where we see it. This sensory congruence is a keystone of our fine hand motor skills.

There are instances, however, where the congruence between hand visual and somatosensory feedbacks is altered, such as when using a microscope or magnifying lenses. In this context, motor performance is disrupted, most probably because the sensorimotor system is fed with conflicting visual and proprioceptive information (Starch, 1910). An interesting support for this hypothesis was provided by Balslev et al. (2004) who showed that a reduction of hand proprioception induced by repetitive transcranial magnetic stimulations (rTMS) of the somatosensory cortex, decreased the detrimental effect of incongruent visual feedback on movement performance. In this novel visuomotor environment, the suppression of somatosensory information would help reduce the sensory conflict, thereby improving motor performance. Note that the results reported by Balslev et al. (2004) are also in line with studies showing that mirror-reversed vision has little impact on the motor performance of patients suffering from a loss of proprioception who trace the contour of a shape (Lajoie et al., 1992; Miall & Cole, 2007).

Previous studies therefore provide clear evidence that processing proprioceptive information is pernicious for controlling movements in the context of a conflict between visual and proprioceptive feedbacks. The question nevertheless remains as to whether the intensity of this conflict is modulated by the possibility/impossibility of seeing the effector from which the conflicting proprioceptive inputs arise. For instance, because the hand muscles are endowed with proprioceptive receptors, the sensory conflict could be enhanced when our hand is visible compared to when we can only see a manipulated tool (e.g., a rod). Indeed, with the sight of the hand, the brain receives visual and somatosensory hand afferents that can be (more or less) directly compared. This context could facilitate detection, and increase the strength, of the sensory mismatch. In this light, it is worth noting that seeing one’s body part has been shown to increase the weight assigned to the somatosensory inputs (Kennett et al., 2001; Longo et al., 2011; Taylor-Clarke et al., 2002, 2004; Zhou & Fuster, 2000). Accordingly, we might expect a greater sensory conflict when tracing the contour of a shape with a hand-held stylus than with a rod, which is devoid of somatosensory attributes.

Here, we tested this prediction by comparing the precision with which healthy human participants traced the contour of a shape with either a stylus (Hand group) or with the tip of a rod (Tool group) in two visual conditions: direct and mirror-reversed vision (i.e., Direct and Mirror conditions, respectively). Based on the hypothesis of a greater sensory conflict when seeing the hand, the tracing performance should be greater for the Tool group than for the Hand group in the Mirror condition. Predictions can also be made regarding the activity of the somatosensory cortex for the Hand and Tool groups when tracing with incongruent visual feedback. Indeed, Bernier et al. (2009) have observed that participants tracing a shape with incongruent visual feedback exhibited a suppression of somatosensory inputs compared to when they were tracing with normal vision. In their study, the somatosensory suppression was evidenced by the decreased evoked potentials within the somatosensory cortex following the electric stimulations of the median nerve at the wrist. Functionally, this suppression of somatosensory information would reduce the sensory conflict (as for the rTMS over the somatosensory cortex, Balslev et al., 2004). Supporting the hypothesis that the sight of the hand increases the visuo-proprioceptive conflict, a gating of somatosensory inputs was not observed by Lebar et al. (2017) when the incongruent hand visual feedback was provided through a digitized dot image (i.e., devoid of somatosensory attributes). In this visual context, Lebar et al. (2017) found a decreased power of beta oscillations (15-25 Hz) in the somatosensory cortex which, on the contrary, reflected greater cortical activity (see Kilavik et al. (2013) for a review on cortical beta oscillations).

Because alpha and beta band powers are respectively considered as being inversely related to the levels of excitability (alpha) and processing (beta) of the somatosensory and visual cortices (Anderson & Ding, 2011; Cheyne et al., 2003; Pfurtscheller & Lopes da Silva, 1999), we predicted that only the Hand group would show greater alpha (8-12 Hz) and beta (15-25 Hz) powers in the somatosensory cortex when tracing with mirror-reversed vision compared to a context with normal vision.

## 2. Method

### 2.1. Participants

Thirty-four volunteers participated to the study. They all had normal or corrected-to-normal vision and were right-handed according to Edinburgh Handedness Inventory (mean laterality score: 77.15 ± 15.4). Informed written consent was obtained before running the experiment. The protocols and procedures were in accordance with the 1964 Declaration of Helsinki and were approved by the CERSTAPS ethic committee. The experiment lasted ~2 hours.

### 2.2. Procedure

The participants were seated in a darkened room in front of an irregular small shape (see Fig. 1) laid on a digitizing tablet. The shape was printed in white on a black background, and was lit by small LEDs directly above the tablet. It was made of 16 thin (0.5 mm) straight lines and 1 curved line whose lengths varied between 8 and 36 mm (total perimeter 36.7 cm). We deliberately chose a complex template (i.e., with many corners) as it has been shown to increase the complexity of the mirror-drawing task (Miall & Cole, 2007). The task consisted of tracing the outline of this shape as precisely as possible with a digitizing stylus (weight 18 grams). The participants of the Hand group (n = 17; 8 women; mean age: 23.7 ± 3.7 years) held the stylus in their right hand. The participants of the Tool group (n = 17) held in their right hand the extremity of a light aluminum rod (40 cm, 17 grams) on the opposite end of which the stylus was firmly fixed. The data of 2 participants had to be discarded because of technical problems. Thus, for the Tool group, the analyses were performed on 15 participants (8 women, mean age: 23.9 ± 2.8 years).

**Figure 1.**
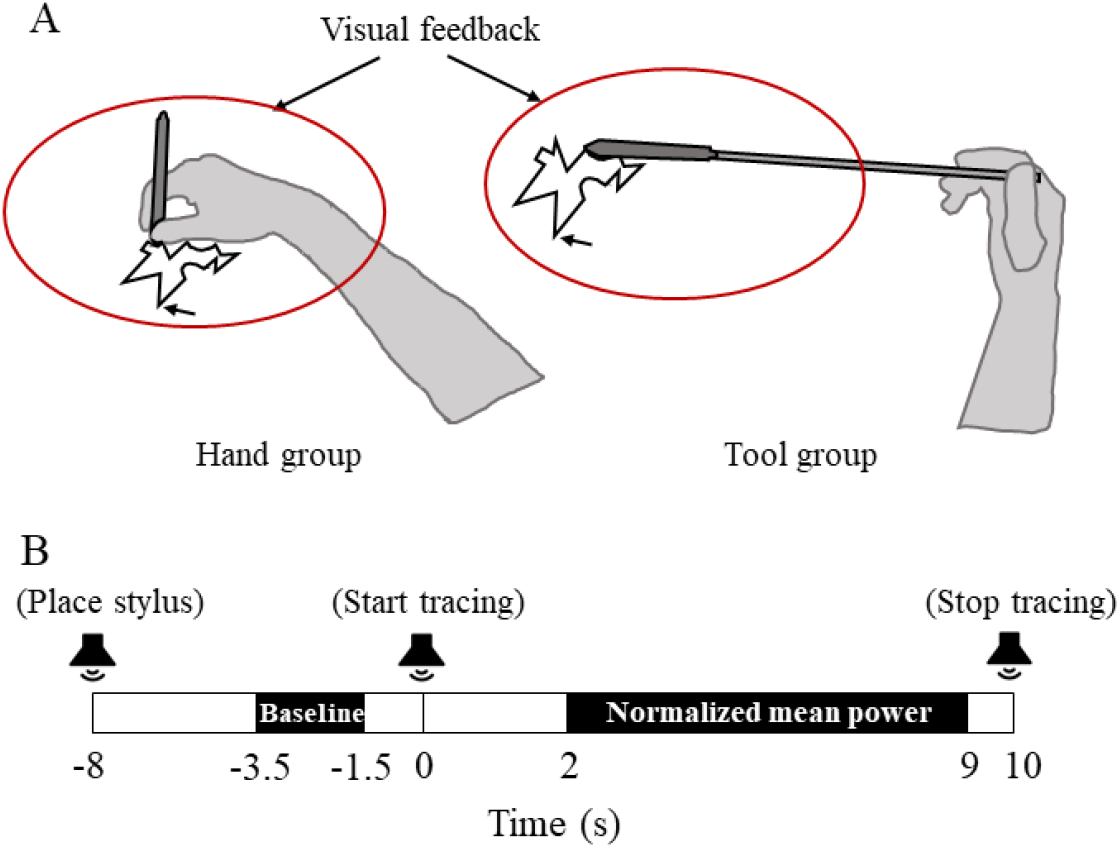
A. Sketches of the visual feedback available during the tracing for the participants of the Hand (left) and Tool (right) groups. The starting position is indicated by the arrows. B. Temporal organization of the trials. For each frequency band (i.e., alpha, beta), the signal computed between 2 s and 9 s after the imperative go signal (i.e., at 0 s) was expressed as a change of power (dB) with respect to a 2-s mean window baseline recorded before the start tracing signal.

Participants of both groups followed the shape in two visual conditions. In the Direct condition, the participants of the Hand group could directly see their hand while participants of the Tool group could see only about the most extreme half of the rod (see Fig. 1A). For this latter group, vision of the arm and the hand was occluded with a black shield. In the Mirror condition, a mirror (Comair Cabinet Executive mirror, diameter 28 cm) was located to the front left of the participant with an inclination of 45 ° relative to the subject’s frontal plane. In this condition, only the hand (Hand group) or the extremity of the rod (Tool group) could be seen through the mirror. For both groups, direct vision of the right upper limb was occluded with a black shield.

Participants of each group performed 40 trials of 18 s duration in both the Direct and Mirror conditions. The temporal organization of every trial is depicted in Fig. 1b. At the beginning of each trial, due to software-related constraints, the tip of the stylus had to be held ~5 cm above the digitizing tablet. For the first trial, all participants held the stylus above the position on the shape indicated by an arrow in Fig. 1a. For the subsequent trials, the participants held the stylus above the position reached at the end of the previous trial. For each trial, with the stylus at these starting positions, the participants sent the verbal message “ready” to the experimenter. Then, on hearing a beep, the participants had to lower the tip of the stylus onto the tablet and to hold the hand and stylus at this position (even if inadvertently the stylus was not on the intended point on the shape). A second beep issued 8 s after the first one served as an imperative signal to start tracing the contour of the shape. A final beep occurring 10 s after the second indicated the end of the trial. All trials were thus composed of a 8 s static phase and of a 10 s dynamic phase. The small size of the shape allowed participants of both groups to perform the tracing using only finger and wrist movements. The participants were instructed that whenever the stylus (or tip of the rod) left the outline of the shape, they should bring it back to the point where it left the shape before continuing the tracing. Participants were required to hold the stylus (Hand group) or the rod (Tool group) with a minimal force and to perform very slow movements. An experimenter demonstrated suitable tracing speeds prior to the experiment and corrective instructions were provided between trials when necessary. Slow movements reduced the muscular activation and the speed of the ocular pursuit which can both contaminate EEG recordings. Offline analyses showed that the mean tracing velocities for the Hand group were 0.54 ± 0.21 cm/s (Direct vision) and 0.47 ± 0.12 cm/s (Mirror vision), and for the Tool group, 0.50 ± 0.11 cm/s (Direct vision) and 0.49 ± 0.11 cm/s (Mirror vision). A 2 x 2 ANOVA did not reveal neither a significant effect of Vision (F_1,31_ = 2.77; p > 0.05) and of Group (F_1,31_ = 0.03; p > 0.05), nor a significant Vision x Group interaction (F_1,31_ = 1.72; p > 0.05).

Our goal was to investigate the effect of seeing one’s hand on the processing of somatosensory information in the context of incongruence between visual and somatosensory feedbacks. Therefore, several elements of the experimental protocol aimed to limit adaptation to the sensory incongruence. The shape had a complex geometry, and the participants had to start their tracing from the position reached in the previous trial in order to avoid an overly repetitive pattern of the layout. The exposure duration to the sensory conflict was only of 6’40’’ (i.e., 40 (trials) × 10 s (dynamic phase duration)). Moreover, after every 5 trials, participants were asked to directly watch their hand moving freely. For reasons of homogeneity between the conditions, this procedure was also followed in the Direct condition.

Participants of both groups were first tested in the Direct condition. Note that contrary to protocols specifically designed to investigate the modification of the internal representation of the body when using tools (e.g., lengthening of the represented arm length, Martel et al., 2016), the present protocol incorporated features to minimize such modifications in the Tool group (e.g., shape positioned in the proximal space, view of the hand moving without the tool every 5 trials).

### 2.3. Data acquisition and processing

#### 2.3.1. Behavior

The X and Y coordinates of the tip of the digitizing stylus were recorded using a Wacom Intuos 4L tablet (spatial resolution of <1mm, 100 Hz recording frequency). The tracing performance was assessed by computing a distance/segment index (referred to as distance ratio) which corresponded to the ratio between the total distance covered by the tip of the stylus and the total length of all drawn segments. The closer this ratio was to 1, the more accurate was the tracing. We also computed the number of reversals in direction when the participants traced the contour of the shape. This was done by calculating and then averaging the number of zero-line crossing in the X and Y velocity of the tracing. The smaller the number of zero-line crossing, the smoother the tracing.

As it can be seen in Fig. 2, both assessments of the tracing performance showed substantial performance deterioration in the Mirror condition for both the Hand and Tool groups. However, performance improved across the first 20 trials before reaching a plateau. In this light, all analyses (i.e., performance, EMG, EEG) were performed using the first 20 EEG artifact-free trials (see below). This series of trial is more likely to better characterize cross-modal conflict between visual and sensorimotor inputs.

**Figure 2.**
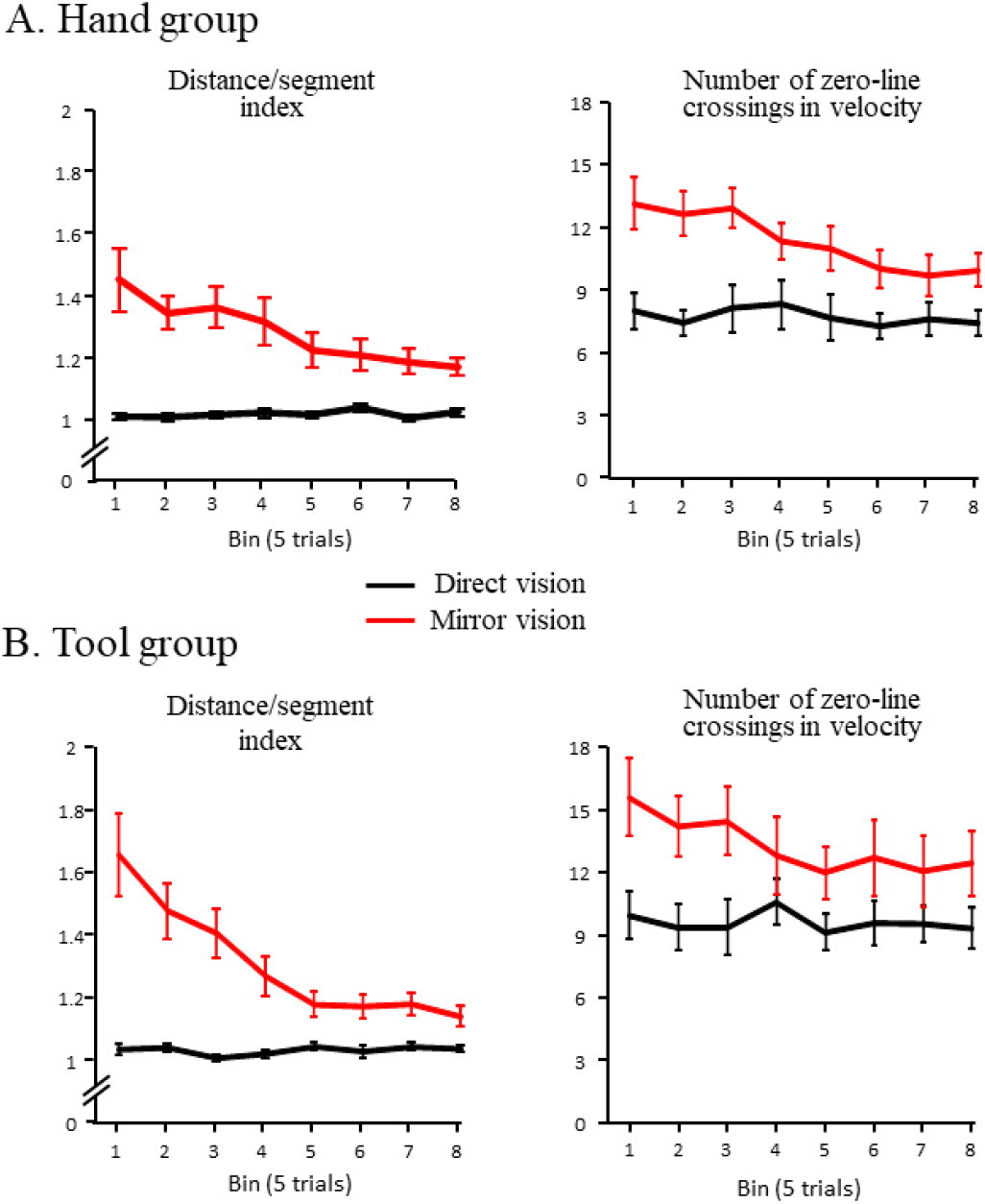
Mean tracing performance over the course of the 40 trials for the Hand group (A) and the Tool group (B). The trials are pooled into 8 bins of 5 consecutive trials. Left panels: The tracing performance is expressed as the average total distance covered by the pen per segment completed in every trial (distance/segment index). Right panels: Number of reversals in direction of the stylus as expressed by the average number of zero-line crossings on the velocity profiles per trial. Error bars: standard error of the mean.

#### 2.3.2. Electromyography (EMG)

The activity of the muscles acting on the wrist and fingers of the right arm was recorded to control for potential large differences of EMG activities between groups and vision conditions. This verification is particularly relevant in the context of the present study because the decrease of proprioception, which is normally observed during movements (Rushton et al., 1981; Seki & Fetz, 2012) is heighten during strong muscle contractions (Staines et al., 1997).

EMG activity was recoded using a Bortec AMT-8 system (Bortec Biomedical, Calgary, Canada; 250 Hz sampling frequency). We recorded the activity of the flexor of the thumb (flexor pollicis brevis) and the first dorsal interosseous muscles, which are both involved in the precision grip. These activities were recorded bipolarly with Ag-AgCl electrodes placed 2 cm apart after cleaning the skin with alcohol. Activity of the flexor and extensor muscles of the wrist was recorded with electrodes placed over the wrist extensor bundle (top of the arm) and over the flexor bundle (bottom of the arm). With this wide configuration, both flexion and extension of the wrist can be recorded with a single pair of electrodes (see Criswell & Cram, 2011, p. 311). An electrode placed above the right epicondyle was used to reference all EMG recordings.

As expected, due to the slow speed of the tracing, the EMG recordings showed tonic activities without clear burst pattern. To compare the EMG activity across groups and conditions, we rectified and integrated the 3 sets of EMG data over both the static phase (−3.5 s to −1.5 s) and the dynamic phase (2 s-9 s) for each valid trial (i.e., without EEG artifact). The integrals (i.e., iEMG) obtained in the dynamic phase were expressed as a percentage of the iEMG obtained in the static phase. Then, we computed the mean % iEMG of the 3 set of EMG data for each group (Hand, Tool) and vision condition (Direct, Mirror).

#### 2.3.3. Electroencephalography (EEG)

EEG activity was recorded continuously using a cap of 64 Ag/AgCl electrodes at a 1024 Hz sampling frequency (ActiveTwo system, Biosemi, Amsterdam, The Netherlands). The activities recorded by electrodes placed near each external canthus, and electrodes placed below and above the left eye were used to detect blinks and saccades. The EEG data were pre-processed using BrainVision Analyzer2 software (Brain Products, Gilching, Germany). EEG signals were referenced against the average of the activities recorded by all electrodes. The effect of ocular artifacts on the EEG recordings, related to blink and saccades, was reduced using the method of Gratton et al. (1983).

For each vision condition, the EEG data were segmented and synchronized with respect to the occurrence of the beep which indicated the beginning of the dynamic phase. Note that due to very slow tracing movements, this segmentation could not be made using kinematic or EMG data within a reasonable temporal margin of error. The recordings were visually inspected and epochs still presenting artifacts were rejected. These trials were replaced by those occurring between the 20^th^ and 27^th^ trials, so that 20 epochs were analyzed for each participant.

We used Brainstorm software to estimate the cortical sources of the EEG signals (Tadel et al., 2011). The inverse problem was resolved using the minimum-norm technique and unconstrained dipole orientations. A boundary element method (symmetric BEM, Gramfort et al., 2010) was used to compute the forward models on the anatomical MRI Colin 27 brain template (15,000 vertices) from the Institut Neurologique de Montréal. We opted for a model with three realistic layers (scalp, inner skull, and outer skull) which yields more accurate solutions compared to a simple three concentric spheres model (Sohrabpour et al., 2015).

Single-trial EEG data were transformed in the time-frequency domain using the Hilbert-filter method. This method is particularly suited for long times-series such as those analyzed in the present study (Cohen, 2014). The analyses of the time frequency distribution were performed in the source space. We extracted the amplitude envelope (i.e., power) of alpha (mean 8-12 Hz, steps of 0.5 Hz) and beta (mean 15-25 Hz, steps of 1 Hz) bands over both the static and dynamic phases of the trials. For each frequency band, the power computed during the dynamic phase was normalized with respect to the static baseline period (−3.5 to −1.5 s) and then averaged, for each group and condition, between all trials over the 2-9 period after the imperative go (“beep”) signal (see Fig. 4). The selected baseline time window was deliberately chosen away from the beep indicating the onset of the static phase, at which time the participants had to lower the stylus on the digitizing tablet (event that was most likely followed by the cognitive appraisal of the stylus landing position). We indistinctly considered increases of alpha and beta band power as a neurophysiological signature of a gating of somatosensory and visual inputs. Decreases of these low and medium frequency bands rather reflecting a facilitation of these sensory inputs. The analyses were limited to the left hemisphere, which was contralateral to the moving (right) hand.

**Figure 3.**
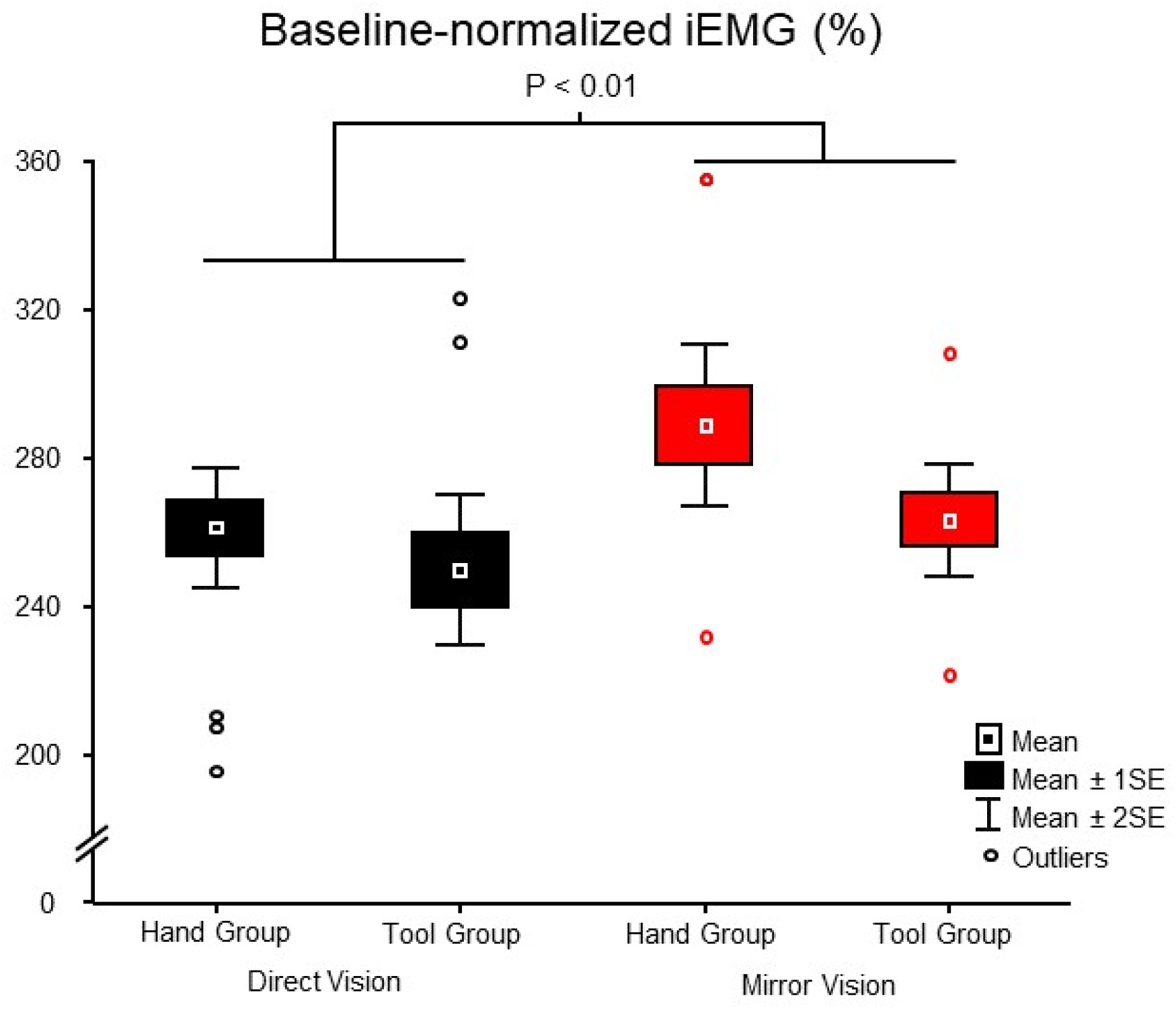
Baseline-normalized iEMG for the Hand and Tool groups computed in the Direct and Mirror conditions.

**Figure 4.**
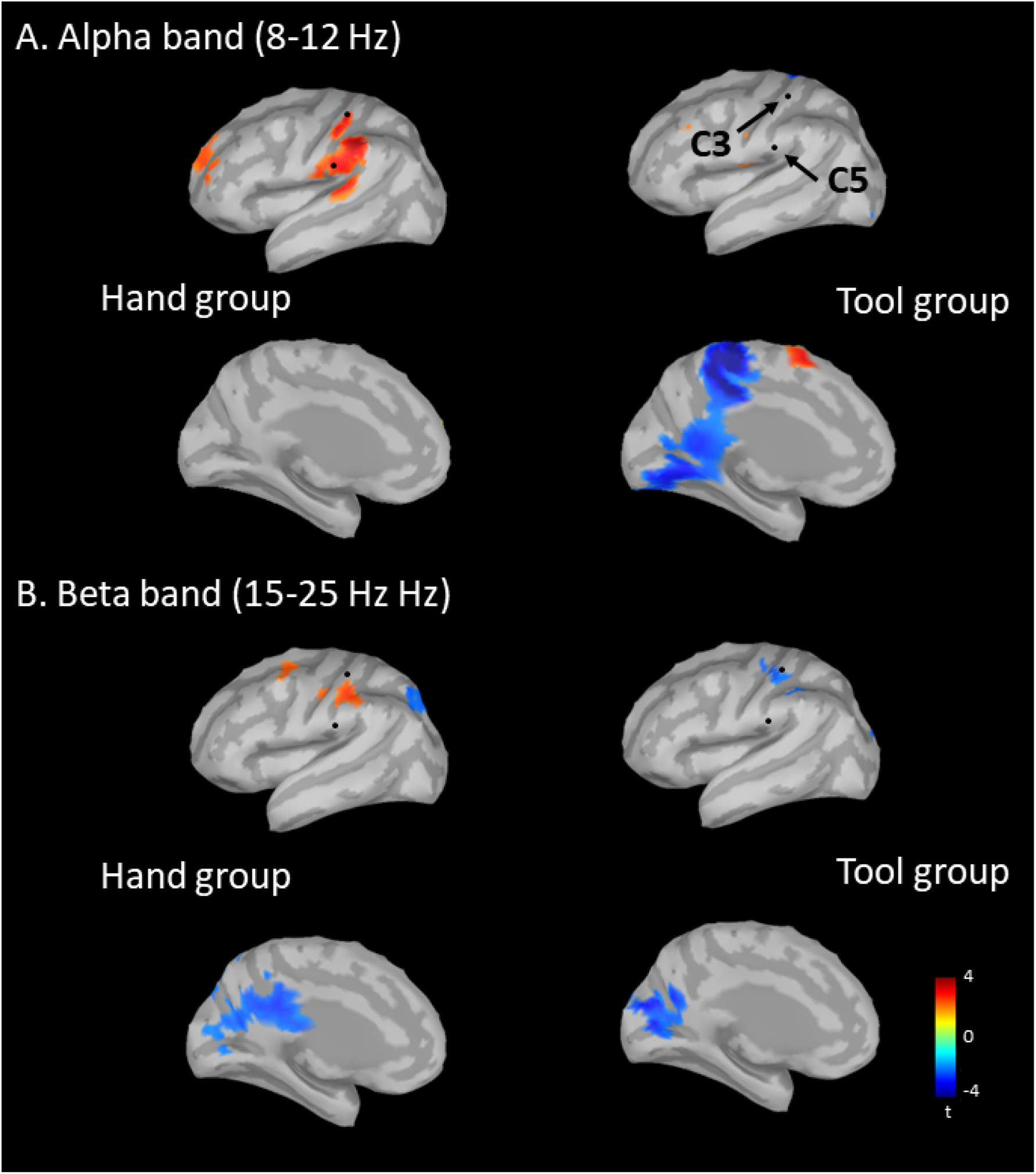
Statistical maps (source space, left hemisphere) of alpha (A) and beta (B) powers resulting from the contrast Mirror vs Normal conditions for both the Hand (left panels) and Tool (right panels) groups. The position of the C3 and C5 electrodes are shown on the side views. These electrodes overlay the left somatosensory cortex (i.e., contralateral to the tracing hand). The signals recorded at these electrodes were used to compare the effect of the visual conditions (i.e., Direct, Mirror) between the Hand and Tool groups (see Fig. 5).

Specific analyses were performed to get insight into the dynamics of the visual feedback-related changes of alpha and beta band powers in the somatosensory and visual cortices. This was done by first identifying from the BEM mesh, and for each participant, the vertex within the somatosensory or visual cortex that exhibited the strongest significant effect (i.e., smallest negative t value or greatest positive t value, see fig. 4) when contrasting the sources of the baseline-normalized alpha and beta band powers estimated in the Direct and Mirror conditions (group analyses, see statistical analyses below). Then, the alpha and beta band powers computed at this vertex in the Mirror condition were extracted from −3.5 s to 9 s, where 0 s indicates the imperative signal to start the tracing movement. Two ways were used to express the time courses of alpha and beta band changes. We computed the mean baseline-normalized power between participants and computed, for each participant, the cumulative integral of the baseline-normalized power. Monotonic increasing or decreasing of the cumulative integral indicates that the increase or decrease of power is preserved throughout the tracing. This computation provides smoother data than the baseline-normalized power and is particularly relevant for appraising the between-participants variability.

The EEG data recorded in the electrode space was also transformed in the time-frequency domain using the Hilbert-filter method. This transformation was performed after applying a spatial filter (surface Laplacian, Perrin et al., 1989; order term of the Legendre polynomial=10, smoothing=1e-5, m=4) thereby increasing the topographical selectivity by filtering out volume-conducted potentials (Law et al., 1993; Nunez & Srinivasan, 2006). Analyzing the spectral content of the EEG signals recorded at C3 and C5 electrodes allowed to directly compare, between the Hand and Tool groups, the effect of tracing with mirror-reversed vision on the alpha and beta band powers over the somatosensory cortex (i.e. the key region for testing the effect vision of the hand on somatosensory processes). Indeed, as shown in Fig. 4, electrodes C3 and C5 respectively overlay the left primary (SI) and secondary (SII, upper bank of the Sylvian fissure) somatosensory cortices.

### 2.4. Statistical analyses

For each Group and Vision conditions, the evolution of the tracing-related variables (i.e., distance/segment index, number of zero speed crossing, iEMG) over the first valid 20 trials was assessed by computing their mean values over 4 bins of 5 consecutive trials. These variables were submitted to separate 2 (Group: Hand, Tool) x 2 (Vision: Direct, Mirror) x 4 (Bin: Bin_1-5_, Bin_6-10_, Bin1_1-15_, Bin_16-20_) analyses of variance (ANOVA), with repeated measurements on the Vision and Bin factors. Significant effects were further analyzed using Newman-Keuls post-hoc tests. The alpha level was set at 0.05 for all statistical contrasts.

For each group, we assessed the effect of the sensory incongruence on the topography and amplitude of the normalized alpha and beta band power by contrasting the sources of alpha and beta band powers estimated in the Direct and Mirror conditions using t-tests (significance threshold p < 0.05, uncorrected).

Finally, to directly compare the effect of the sensory incongruence on somatosensory alpha and beta band powers between the Hand and Tool groups, we subtracted for both the C3 and C5 electrodes and for all participants of each group, the normalized power computed in the Mirror condition from the normalized power computed in the Direct condition. The differences (hereafter referred to as ΔMirror-Direct) were submitted to independent T-tests (significance threshold p < 0.05).

## 3. Results

### 3.1. Tracing performance

The evolution of the distance/segment index and of the number of zero speed crossing throughout the 40 trials are shown in Fig. 2 for both the Hand and Tool groups. Overall, the participants of both groups accurately traced the shape with Direct vision but substantially decreased their tracing accuracy with mirror-reversed vision. Figure 2 shows improvement in tracing performance over the first 20 trials before reaching a relative stable plateau, suggesting that the sensory conflict was perceived greater in the first half of the trials. Because our main goal was to compare the response of the somatosensory cortex when tracing a shape in the context of a visuo-proprioceptive conflict, all behavioral and electrophysiological analyses presented below pertained to the first 20 trials (see methods for exceptions).

The distance/segment index was significantly greater in the Mirror (mean: 1.41 ± 0.48) than in the Direct (mean: 1.02 ± 0.04) conditions (main effect of Vision: F_1,31_ = 63.65; p < 0.001; η^2^ = 0.68). For this variable, the ANOVA did not reveal a significant effect of Group (F_1,31_ = 0.78; p > 0.05), but revealed a significant Vision x Bin interaction (F_1,31_ = 6.89; p < 0.001; η^2^ = 0.19). Post-hoc analyses confirmed the decrease of the distance ratio over the trials with mirror-reversed vision, but more importantly, they showed that the distance ratio computed in the last series of 5 trials (i.e., bin no. 4) was still significantly greater than the distance ratio computed in all bins of the Direct condition (all ps < 0.05).

The number of zero speed crossing was also significantly greater in the Mirror (mean: 13.44 ± 5.04) than in the Direct (8.92 ± 3.68) conditions (main effect of Vision: F_1,31_ = 44.04; p < 0.001). For this variable, the ANOVA did not reveal neither a significant effect Group (F_1,31_ = 1.51; p > 0.05), nor a significant Vision x Bin interaction (F_1,31_ = 2.38; p > 0.05).

### 3.2. EMG recordings

Figure 3 shows the iEMG, computed from the recordings of the forearm and hand muscles during the tracing, normalized to the iEMG computed before starting the tracings. The figure shows that the iEMG was ~200-300% greater during the tracing compared the static period. The ANOVA revealed that the normalized iEMG was significantly greater in the Mirror condition (mean: 278% ± 40) than in the Direct condition (mean: 257% ± 34) (F_1,31_ = 11.05; p < 0.005; η^2^ = 0.29). However, the effect of Group (F_1,31_ = 2.26; p > 0.05), the interaction between Vision and Group (F_1,31_ = 1.31; p > 0.05) and the interaction between Vision and Bin (F_1,31_ = 0.12; p > 0.05) were not significant. Therefore, if different spectral contents of cortical neural oscillations were to be found between the Hand and Tool groups, they would unlikely result from different muscular activities (see Staines et al., 1997 for the effect motor contractions amplitude on the gating of somatosensory inputs). The increased hand muscle activities observed with mirror-reversed vision could be due to the greater number of reversals in direction when tracing the contour of the shape with incongruent vision (Fig. 3).

### 3.3. EEG data

Figure 4 shows the statistical maps of alpha and beta band power resulting from the contrast Mirror vs Direct conditions for both the Hand and Tool groups. Warm colors indicate that alpha and beta band powers were significantly greater in the Mirror condition than in the Direct condition. If observed in sensory areas, warm colors would therefore reflect a relative decrease in weight assigned to the inputs pertaining to these areas when tracing with mirror-reversed vision. Cold colors indicate the opposite pattern. Remarkably, the significant differences resulting from the contrasts Mirror vs Direct conditions were largely circumscribed to the somatosensory and visual areas for the Hand group, and to visual areas for the Tool group.

#### 3.3.1. EEG data: Somatosensory cortex

For the Hand group, alpha band power was significantly greater when tracing with mirror-reversed vision in areas identified by the source analyses as the primary (SI) and the secondary (SII, i.e. upper bank of the Sylvian fissure) somatosensory cortices (Fig. 4a). Beta band power was also significantly greater with incongruent visual feedback in SI (Fig. 4b). For the Tool group, alpha and beta band powers computed in the somatosensory cortex were strikingly alike between the Mirror and Direct conditions. The statistical map only revealed a significantly smaller alpha band power in a small area of SI (Fig. 4a).

Alpha and beta band powers recorded at C3 and C5 electrodes were also compared between Groups and Vision conditions over the same time windows as the analyses in the source space. These electrodes overlay the left postcentral region (Koessler et al., 2009, see also Fig. 4) which was contralateral to the tracing hand. T-tests revealed that the ΔMirror-Direct beta (t(30) = 3.01; p < 0.01; d = 0.95) and the ΔMirror-Direct alpha (t(30) = 2.50; p < 0.01; d = 0.83) significantly differed between groups at electrode C3 and C5, respectively (Fig. 5). Importantly, for the Hand group, the ΔMirror-Direct beta value (electrode C3) was positive (mean = 9.87 ± 12.77) and was significantly different from 0 (comparison to a standard (i.e., 0); p < 0.01). Likewise, for the Hand group, the ΔMirror-Direct alpha value (electrode C5) was positive (mean = 9.70 ± 18.79) and also significantly differed from 0 (p < 0.05). However, the ΔMirror-Direct alpha (C3) and the ΔMirror-Direct beta (C5) did not significantly differ between groups (t(30) = 0.93; p > 0.05 and t(30) = 1.23; p > 0.05, for C3 and C5, respectively). For the Tool group, the ΔMirror-Direct alpha and beta bands computed at electrodes C3 and C5 did not significantly differ from zero (ps>0.05).

**Figure 5.**
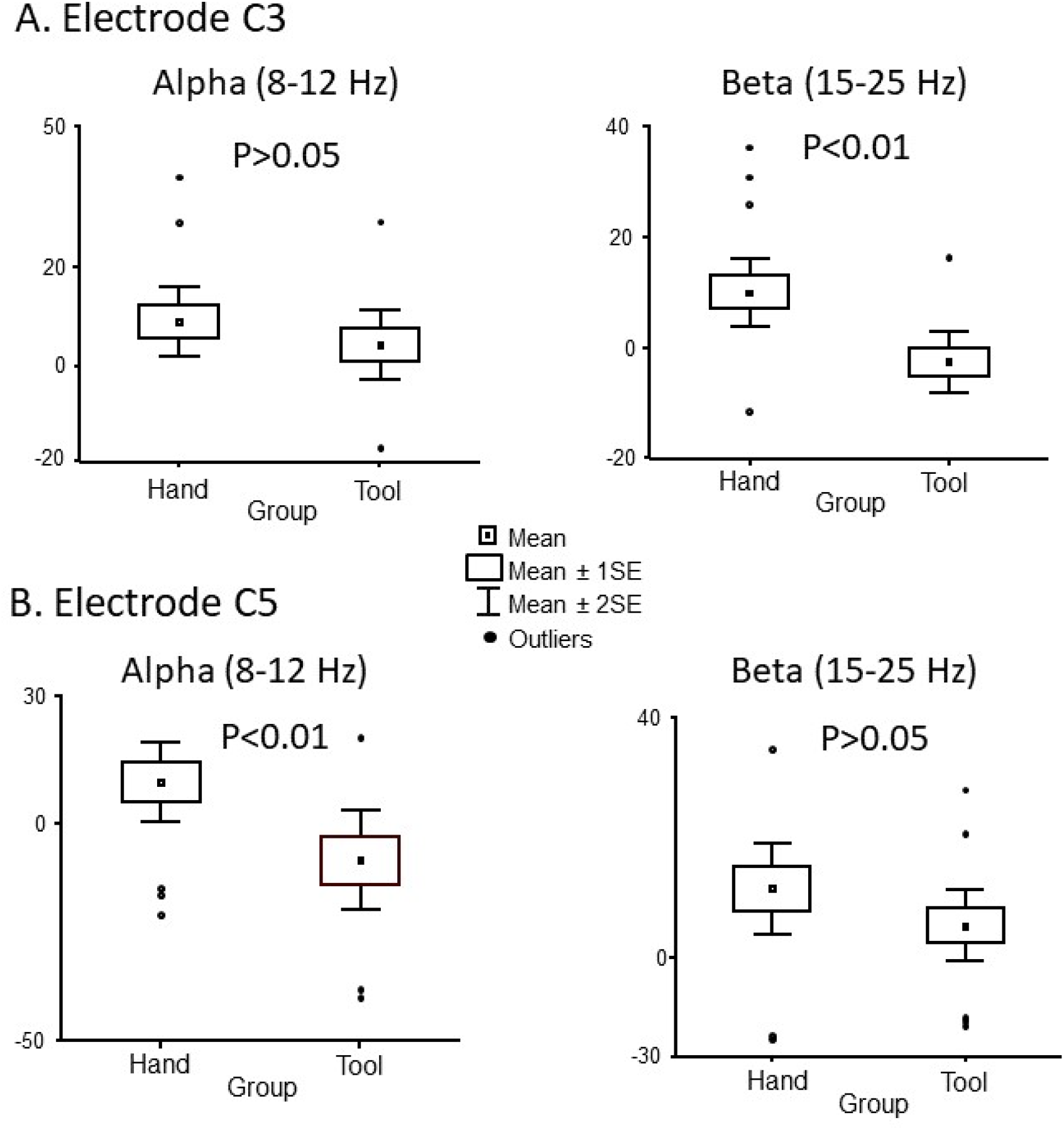
Comparison between the ΔMirror-Normal (alpha and beta, expressed in signal units^2^/Hz) computed at electrode C3 (A) and electrode C5 (B) for the Hand and Tool groups. These electrodes overlay the left sensorimotor cortex (see Fig. 4). The significant effect of Group was preserved at electrode C3 (t(25) = 3.25; p < 0.005) and at electrode C5 (t(24) = 3.18; p < 0.005) when performing the tests after removing the outliers.

#### 3.3.2. EEG data: Visual cortex

The power within alpha and beta bands computed in the medial visual cortex was also altered when tracing with mirror vision. In contrast to what was observed in the somatosensory cortex, the bias in the visual feedback led to decreases in alpha and beta band powers in visual areas (Fig. 4). This suggests a facilitation of visual feedback with mirror-reversed vision. However, the effect of the incongruent visual feedback on the neural oscillations appeared more pronounced for the Tool group than for the Hand group. Indeed, the statistical maps showed significant smaller power in the Mirror condition in regions estimated by sources analyses as the lingual gyrus (alpha), the medial parietal cortices (alpha) and the cuneus (beta). For the Hand group, the statistical maps only revealed significantly smaller beta band power in the cuneus (Fig. 4). Note that because the effects of mirror-reversed vision occurred in the medial visual cortex, the ΔMirror-Direct alpha and beta band powers could not be computed in the electrode space.

The contrast Mirror vs Direct also revealed smaller alpha band power in the Mirror condition for the Tool group in a region identified as the anterior precuneus cortex.

### 3.4. EEG data: dynamics of the changes of alpha and beta band powers

Figure 6 provides an estimate of the dynamics of the increased in alpha and beta band powers when the participants of the Hand and Tool groups traced the shape with mirror-reversed feedback. Band powers were extracted from vertices within areas showing significant contrasts between the Mirror and Direct conditions (i.e., SI, SII, cuneus, lingual gyrus, see Fig. 4). The figure shows that the increased in power observed in the somatosensory cortex (results obtained only for the Hand group) was more consistent in SII (alpha) than in SI (beta). Indeed, 14 out of 17 participants showed an increase of alpha band power in SII when they traced the shape in the Mirror condition while 10 participants showed an increased beta in SI.

**Figure 6.**
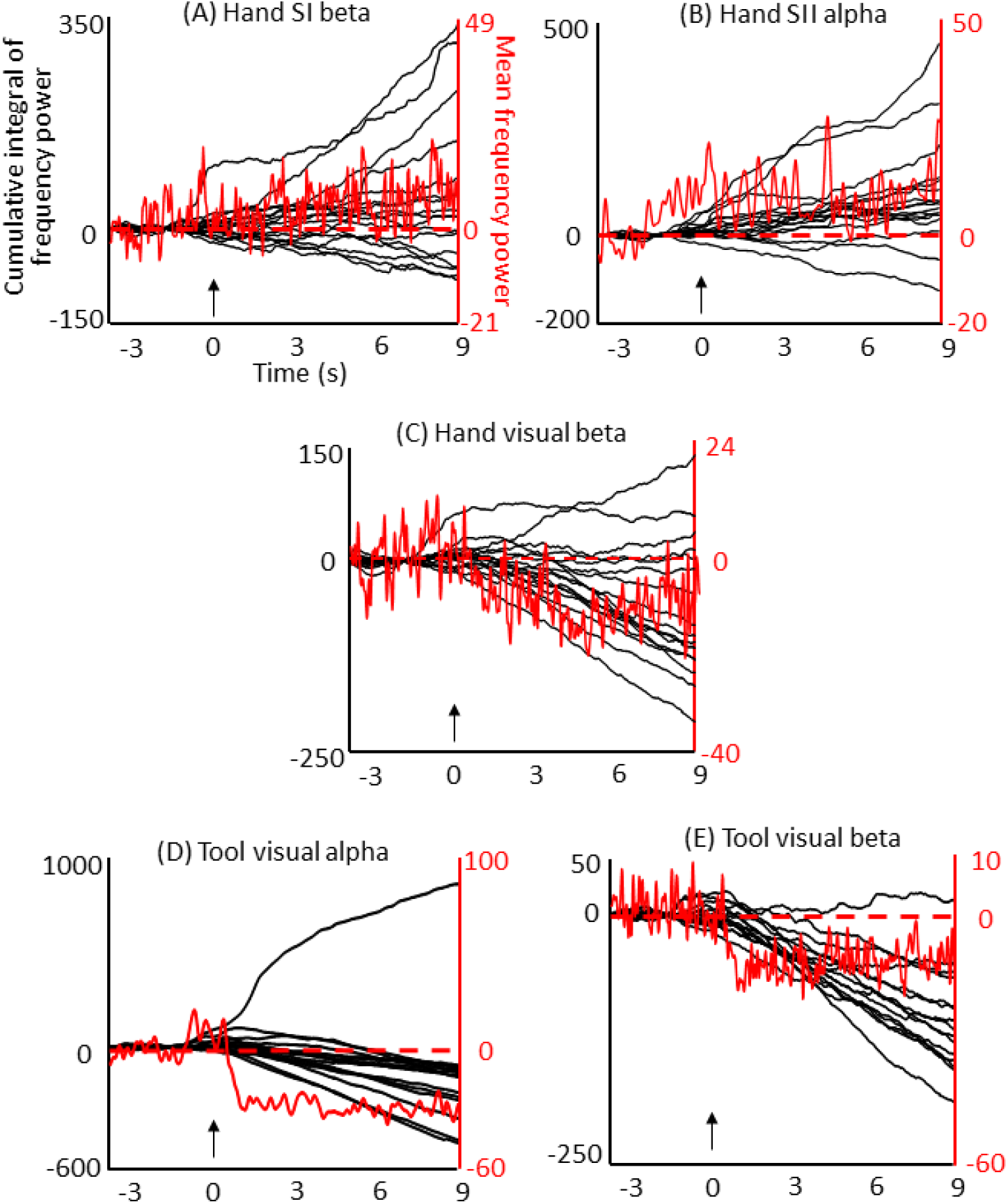
Time course of the baseline-normalized alpha and beta powers. For each participant, the powers (signal units^2^/Hz) were extracted from vertices within areas showing, for either the Hand or the Tool groups, significant contrasts between the Mirror and Normal conditions (see Fig. 4). The red traces represent the between-participants mean powers. The black traces represent the cumulative integral of the baseline-normalized power computed for each participant. The arrows indicate the start tracing signal.

On the other hand, the decrease in alpha and beta band powers observed in the visual cortex, when tracing with mirror-reversed vision, was more robust in the Tool group compared to the Hand group; the power decreased in all participants except one. Remarkably, the participant in the Tool group showing a larger increase in alpha band power during mirror vision, also had the greatest number of zero-line crossings in tracing velocity (i.e., worst tracing performer).

Together, these results are consistent with those issued from the statistical maps (Fig. 4) that showed i) for the Hand group, a greater cluster exhibiting significant increase in alpha band power sources localized in SII, and ii) for the Tool group, greater clusters exhibiting significant decreases of alpha and beta band powers in the visual cortex.

## 4. Discussion

We tested the hypothesis that the conflict between visual and arm proprioceptive inputs, when tracing the contour of a shape with mirror-reversed vision, is greater when participants see their hand during tracing. Contrasting the spectral content of the cortical oscillatory activity in conditions with and without incongruent visual feedback (respectively Mirror and Direct conditions), we observed increases of alpha and beta band powers in the somatosensory cortex when participants had vision of their hand when tracing with mirror vision (Hand group). In contrast, for participants tracing with the tip of a rod (i.e., without hand visual feedback, Tool group), alpha and beta band powers in the somatosensory cortex did not significantly differ between the Direct and Mirror conditions. There is a consensus that increases in alpha and beta band powers respectively correspond to a decrease in cortical excitability and processing (Anderson & Ding, 2011; Cheyne et al., 2003; Kilavik et al., 2013; Pfurtscheller & Lopes da Silva, 1999). In this light, the changes of alpha and beta band powers observed in the somatosensory cortex imply a suppression of arm somatosensory information. The fact that only the participants of the Hand group showed a gating of arm somatosensory inputs with mirror vision suggests that seeing the hand enhanced the visuo-proprioceptive conflict. Altered visual feedback, however, deteriorated tracing performance similarly in both the Hand and Tool groups. The results showed by the participants of the Tool group suggest that their altered performance with mirror vision essentially stemmed from a sensory-motor conflict (rather than from a visuo-proprioceptive conflict, see below).

Moving our arm or an object when seen through a mirror creates a mismatch between the movement-related information carried by the visual and proprioceptive systems. Conceptually, this mismatch prevailed in the present experiment when the participants of both the Hand and Tool groups traced the shape with mirror vision. However, only the participants of the Hand group showed a suppression of somatosensory information (i.e., greater alpha and beta band powers in the Mirror condition). Functionally, the dynamic suppression of somatosensory information when performing goal-directed movement under incongruent visual inputs is thought to reduce the sensory conflict (Bernier et al., 2009; Goldenkoff et al., 2021). Within this framework, our results are then compatible with two non-mutually exclusive scenarios. One in which vision of the hand would enhance arm somatosensory information, thereby increasing the sensory conflict. This would be consistent with psychophysical studies showing enhanced processing of somatosensory information (from extraocular, neck and arm muscles) with visual feedback (Becker & Saglam, 2001; Blouin et al., 2002; Kennett et al., 2001; Longo et al., 2011; Taylor-Clarke et al., 2002, 2004; Zhou & Fuster, 2000). It would also be compatible with the greater sensitivity of the somatosensory cortex to peripheral somatosensory inputs reported in previous studies when the stimulated body area can be seen (Forster & Eimer, 2005; Sambo et al., 2009; Taylor-Clarke et al., 2002). Another possibility is that the inter-sensory conflict increased for the Hand group because the source of the conflicting somatosensory inputs (i.e., the hand) could be seen, contrary to the Tool group. According to this hypothesis, the view of the hand would allow a more direct comparison between the visual and somatosensory mapping of the hand, and therefore a better detection of a sensory mismatch when controlling movements with incongruent visual feedback.

Our results point to an automatic covert processing of arm proprioceptive inputs induced by vision of the hand. In normal visual condition, this covert processing might contribute to the high quality of our broad manual motor repertoire. In conditions with incongruent visual feedback, it would impair movement performance, thereby prompting the brain to decrease the weight of proprioception during the visual and somatosensory feedbacks integration. The fact that the participants of the Tool group did not show significant modulation of somatosensory alpha and beta powers when tracing with mirror vision suggests that vision of a self-moved tool does not enable such covert processing of proprioceptive information. In the present study, we did not control for change of the internal representation of the body when using tools (e.g., lengthening of the represented arm length, Cardinali et al., 2009; Sposito et al., 2012; see Martel et al., 2016 for a review). However, our experiment was designed to minimize such modifications (e.g., participants viewed their hand moving without the tool every 5 trials). Further studies are needed to determine whether the view of the tool also leads to down-weighting of proprioception in the somatosensory cortex when the tool is incorporated into body representations.

The present results revealed that the dynamic control exerted by the brain over arm somatosensory information mainly occurred in SII, which is an important hub for processing somatosensory information (Steinmetz et al., 2000). Our findings are then consistent with studies showing greater attention-related processes in SII than in SI (Chapman & Meftah, 2005; Nelson et al., 2004). Importantly, SII is thought to contribute to the integration of proprioceptive inputs for the online motor control (Eickhoff et al., 2010; Hinkley et al., 2007). The sensory gating observed in SII areas then likely decreased the weight assigned to arm proprioceptive inputs when controlling movements with incongruent visual and proprioceptive feedbacks.

Although occurring outside our pre-defined region of interest (i.e., somatosensory area), we found significant decreases of alpha and beta powers in the occipital cortex when participants traced the shape with mirror-reversed vision. The effect of the incongruent visual feedback on the activity of the occipital cortex was therefore opposed to the effect observed in the somatosensory cortex for the Hand group (i.e., increased alpha and beta powers). The decrease in occipital alpha and beta band powers is consistent with a facilitation of visual inputs when performing movement under visuo-proprioceptive incongruence. This change in occipital alpha and beta powers corroborates brain imaging studies (e.g., EEG, fMRI) reporting increased activity in the occipital lobe when performing movements under discrepant visual feedback (Lebar et al., 2015; Limanowski et al., 2017, 2020).

The changes in alpha and beta band powers observed in the occipital were more robust for the Tool than for the Hand groups. This observation suggests that seeing a self-moved tool under incongruent visual feedback is a favorable context to create a visual attentional set, which increases visual brain activity (see Limanowski & Friston, 2019; Limanowski, 2022). On the other hand, for the Tool group, the shift of attention away from arm proprioception (and perhaps away from hand working space), and the absence of covert processing of arm proprioceptive inputs in the absence of hand visual feedback, might have reduced the weight of arm proprioceptive inputs when tracing the shape with normal visual feedback. Viewed from this perspective, there would be no functional necessity to further downregulate arm somatosensory inputs when tracing with incongruent visual feedback. This could explain why, contrary to the Hand group, the Tool group showed similar somatosensory alpha and beta band powers between the Mirror and Direct conditions. Therefore, the present results could reconcile the apparent discrepancy between the suppression of somatosensory inputs reported by Bernier et al. (2009) when participants traced the contour of a shape while seeing their hand through an inclined mirror (as in the present study) and the reduction of somatosensory beta band power (i.e., increased processing) reported by Lebar et al. (2017) when the incongruent hand visual feedback was provided using a digitized dot image.

Our source analyses estimated the cuneus (for the Hand and Tool groups) and the lingual gyrus (for the Tool group) as the origin of the occipital decrease of alpha and beta band powers in the Mirror condition. These medial visual areas are known to encode space in an allocentric frame of reference (Chen et al., 2014; Committeri et al., 2004; Ruotolo et al., 2019). In this frame of reference, the body (including the hand) and the objects of the environment would be encoded relative to each other within a retinal map (i.e., object-based coding of space) (Burgess et al., 2004; Galati et al., 2000; Paillard, 1987). Such visual representation of space would be largely independent of somatosensory inputs (Ambrosini et al., 2012; Blouin et al., 1993; Medendorp et al., 2008). Accordingly, our results suggest that controlling the motion of the hand or of a tool with incongruent visual feedback enables the use of an allocentric reference frame. The fact that the Tool group showed stronger between-subjects consistency regarding the decreased alpha and beta band powers in the medial visual cortex implies that the manipulated tool was selectively encoded with an object-based frame of reference. The observation that the only participant of the Tool group who showed a strong increase in alpha band power in the Mirror condition was the worst tracing performer provides evidence that this frame of reference was more relevant for controlling arm movements in this novel visuomotor environment than somatosensory-based egocentric reference frames. Moreover, the finding of both increase and decrease of visual beta band power when the participants of the Hand group traced with mirror vision supports the suggestion that the selection of the frames of reference is subject and context dependent (Bernier & Grafton, 2010; Bridgeman, 1991; Byrne & Henriques, 2013). The enhanced object-based coding of space for the Tool group in the Mirror condition is also supported by the decreased alpha band power observed in the anterior precuneus with mirror vision. Indeed, this medial area of the parietal cortex has been shown to selectively encode the motor goal in visual coordinates (Bernier & Grafton, 2010).

We reasoned that because the tool is devoid of somatosensory attributes, the visuo-proprioceptive conflict should be less perceived for the Tool group. Accordingly, we predicted better performance for the Tool than for the Hand groups in conditions with incongruent visual feedback. Behavioral analyses rather revealed that the tracing performance of both groups was similarly impaired with mirror vision. A likely explanation is that the performance degradation showed by the Tool group mainly resulted from a sensorimotor conflict (rather than from a visuo-proprioceptive conflict). During visually-guided movements, this conflict would result from the incongruence between the actual visual feedback and the predicted visual feedback issued from the motor commands (Brun et al., 2020; Miall & Cole, 2007; Shadmehr et al., 2010). Similar conflict could have emerged between the predicted and the actual somatosensory feedbacks. In our study, the hand motor commands when manipulating the tool might have enabled these sensory predictions. Most likely, the visuomotor conflict also degraded the tracing performance of the Hand group. However, the fact that for the Tool group, the incongruence between visual and somatosensory feedbacks had no significant impact on the somatosensory alpha and beta band powers suggests that the visuomotor conflict had only negligible effect on the activity of the somatosensory cortex.

## 5. Conclusion

We found that the control of tracing movement under incongruent visual and somatosensory information was associated with an increased alpha (8-12 Hz) and beta (15-25 Hz) band powers in the somatosensory cortex if participants had visual feedback of their hand. This modulation of alpha and beta activities, which suggested reduced proprioception, was not found if participants traced the shape with the tip of a rod without seeing their hand. Taken together, our findings are in line with a covert processing of arm somatosensory information induced by vision of the hand. This convert processing would have a detrimental effect on movements that are controlled under incongruent visual and proprioceptive feedbacks, and would prompt the brain to exert a control over somatosensory information. Our results suggest that the processing of arm somatosensory inputs during the control of goal-directed hand movements differs largely between conditions where hand visual feedback is available and conditions where the hand cannot be seen. This could explain results from previous studies (e.g., Clower & Boussaoud, 2000; Norris et al., 2001) showing that the sensorimotor adaptation to prismatic displacement is greater when the participants can see their hand than when the participants see their hand in a more abstract form (e.g., digitized dot, video).

## Acknowledgments

The authors thank Marie Fabre, Aurélie Grandjean and Chloé Sutter for their help at various stages of this research project.

## Credit Authorship Contribution Statement

Benjamin Mathieu: Conceptualization, Methodology, Investigating, Data analyses, Writing, Reviewing & Editing. Antonin Abillama: Conceptualization, Methodology, Investigating, Data analyses. Simon Moré: Sofware. Catherine Mercier: Conceptualization, Reviewing & Editing. Martin Simoneau: Conceptualization, Reviewing & Editing. Jérémy Danna: Software, Reviewing & Editing. Laurence Mouchnino: Conceptualization, Methodology, Reviewing & Editing. Jean Blouin: Conceptualization, Methodology, Investigating, Data analyses, Writing, reviewing & editing.

